## Supplementary material for "Disease progression modelling reveals heterogeneity in trajectories of Lewy-type α-synuclein pathology"

**SUPPLEMENTARY TABLES**

**Table S1. Demographics of the SuStaIn modelling cohort.**

|  | **SuStaIn modelling cohort** |
| --- | --- |
| ***N* (%)** | 814 |
| **Age at death (years)** | 81.8 (7.7) |
| **Female, n (%)** | 317 (38.9) |
| **Education, years**^a^ | 14.9 (2.8) |
| ***APOE* ε4 carrier, n (%)**^b^ | 340 (41.8) |
| **MMSE score**^c^ | 17.0 (9.6) |
| **△t MMSE and death (months)** | 21.8 (21.5) |
| **Clinicopathological diagnosis** |  |
| **AD, n (%)** | 285 (35.0) |
| **PD, n (%)** | 168 (20.6) |
| **DLB, n (%)** | 19 (2.3) |
| **Mixed AD and PD, n (%)** | 78 (9.6) |
| **Mixed AD and DLB, n (%)** | 141 (17.3) |
| **ILDB, n (%)** | 90 (11.1) |
| **Other**^d^**, n (%)** | 33 (4.1) |
| **Total Lewy body density**^e^ | 18.1 (11.6) |
| **Total plaque load**^f^ | 9.74 (5.65) |
| **Total neurofibrillary load**^g^ | 8.78 (4.7) |
| ***Post-mortem* interval (hours)** | 5.43 (9.7) |

Data are shown as mean (standard deviation) unless otherwise specified.

^a^ *n*=659.

^b^ *n*=807.

^c^ *n*=672.

^d^ Other indicates the absence of an AD or Lewy body disorder diagnosis.

^e^ Sum of the regional density scores (range 0-40), *n=*701.

^f^ *n*=804.

^g^ *n*=801.

AD = Alzheimer’s disease; DLB = dementia with Lewy bodies; ILDB = incidental Lewy body disease; PD = Parkinson’s disease.

**Table S2. Frequency of other clinicopathologic diagnoses in the SuStaIn modelling cohort.**

|  | **SuStaIn modelling cohort with any other clinicopathological diagnosis (*n*=33)** |
| --- | --- |
| **Argyrophilic grain dementia** | 13 (39.4) |
| **Progressive supranuclear palsy** | 11 (33.3) |
| **Cerebral amyloid angiopathy** | 9 (27.3) |
| **Vascular dementia** | 7 (21.2) |
| **Hippocampal sclerosis** | 5 (15.2) |
| **FTLD-TDP**^a^ | 4 (12.1) |
| **Corticobasal degeneration** | 2 (6.1) |
| **Pick’s disease** | 2 (6.1) |
| **Dementia lacking distinctive histology** | 1 (3.0) |
| **Motor neuron disease** | 1 (3.0) |
| **Huntington’s disease** | 1 (3.0) |

Other is defined as the absence of a Lewy body disease or Alzheimer’s disease diagnosis. Data are shown as number (percentage).

^a^ not assessed for *n*=15 (45.5%).

FTLD-TDP = Frontotemporal lobar degeneration with TPD-43-immunoreactive pathology.

**Table S3. Lewy type α-synuclein scores of subjects assigned to SuStaIn stage 0.**

| **OBT** | **Medulla** | **Pons** | **Substantia nigra** | **Amygdala** | **Transentorhinal cortex** | **Cingulate cortex** | **Temporal cortex** | **Frontal cortex** | **Parietal cortex** |
| --- | --- | --- | --- | --- | --- | --- | --- | --- | --- |
| 0 | 1 | 0 | 0 | 1 | 0 | 0 | 0 | 0 | 0 |
| 0 | 0 | 0 | 0 | 1 | 0 | 0 | 0 | 0 | 0 |
| 0 | 1 | 0 | 0 | 0 | 0 | 0 | 0 | 0 | 0 |
| 0 | 1 | 0 | 0 | 0 | 0 | 0 | 0 | 0 | 0 |
| 0 | 1 | 0 | 0 | 0 | 0 | 0 | 0 | 0 | NA |
| 0 | 0 | 0 | 1 | 1 | 0 | 0 | 0 | 0 | 0 |
| 0 | 0 | 0 | 0 | 1 | 0 | 1 | 0 | 0 | 0 |
| 0 | 1 | 0 | 0 | 0 | 0 | 0 | 0 | 0 | 0 |
| 0 | 1 | 0 | 0 | 0 | 0 | 0 | 0 | 0 | 0 |
| 0 | 0 | 0 | 0 | 2 | 0 | 0 | 0 | 0 | 0 |
| NA | 0 | 1 | NA | 0 | 0 | 0 | 0 | 0 | 0 |
| 0 | 0 | 0 | 0 | 1 | 0 | 0 | 1 | 0 | 0 |
| 0 | 0 | 0 | NA | 1 | 1 | 0 | 0 | 0 | 0 |

Description in terms of Lewy type α-synuclein density scores of subjects assigned to SuStaIn stage 0. Scores range from 0 to 4, with higher scores indicating more severe pathology.

**Table S4. Neuropathological density scores converted to score probabilities.**

|  |  | **Score Probability** | | | | |
| --- | --- | --- | --- | --- | --- | --- |
|  |  | 0 | 1 | 2 | 3 | 4 |
| **Neuropathological density score** | 0 | 0.88 | 0.12 | 2.95 x 10^-4^ | 1.34 x 10^-8^ | 1.12 x 10^-14^ |
|  | 1 | 0.11 | 0.79 | 0.11 | 2.64 x 10^-4^ | 1.20 x 10^-8^ |
|  | 2 | 2.64 x 10^-4^ | 0.11 | 0.79 | 0.11 | 2.64 x 10^-4^ |
|  | 3 | 1.20 x 10^-8^ | 2.64 x 10^-4^ | 0.11 | 0.79 | 0.11 |
|  | 4 | 1.12 x 10^-14^ | 1.34 x 10^-8^ | 2.95 x 10^-4^ | 0.12 | 0.88 |
|  | Missing | 0.20 | 0.20 | 0.20 | 0.20 | 0.20 |

**Figure S1. Model fit for up to five SuStaIn subtypes**

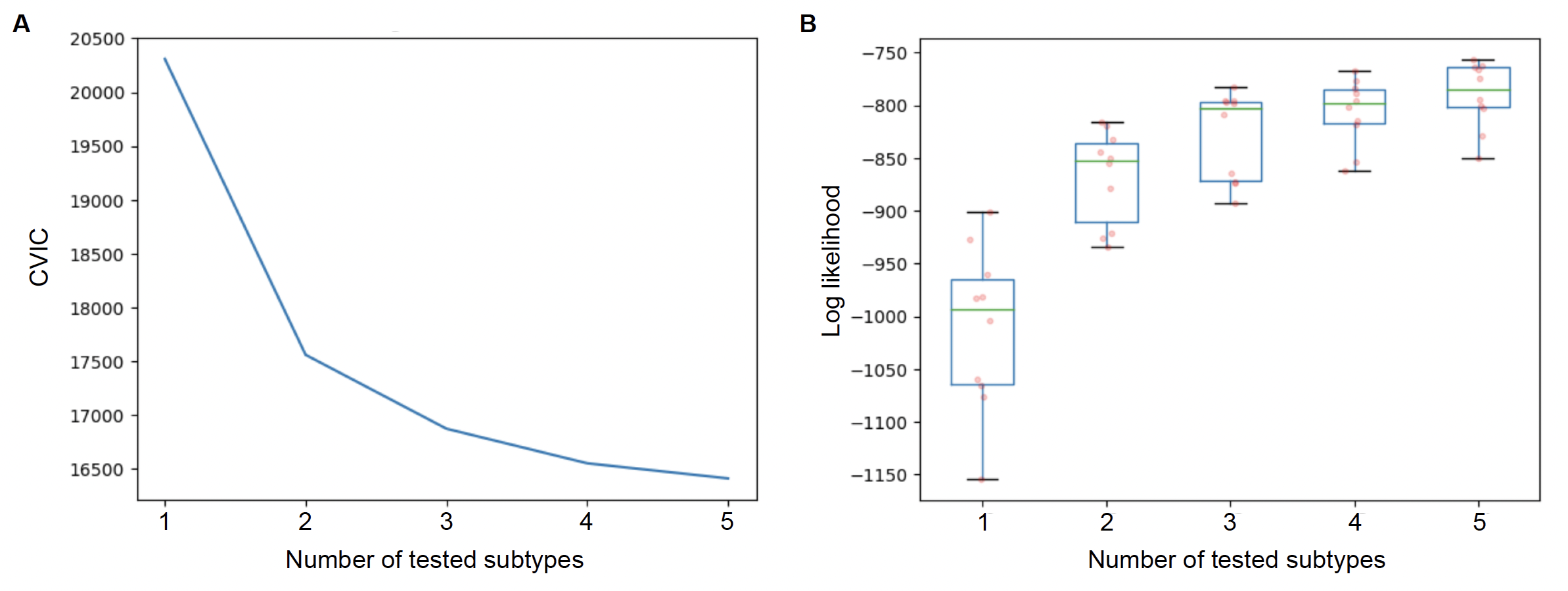

Model fit for SuStaIn models with varying number of subtypes. **A** Cross-validation information criterion (CVIC) and **B** log-likelihood across 10-folds of cross-validation are shown for 1-5 subtypes. Substantial improved model fit can be appreciated from one to two and two to three subtypes, but not for more subtypes. Therefore, three subtypes were selected.

**Figure S2 Output of the SuStaIn model applied to complete data only (*N*=701)**

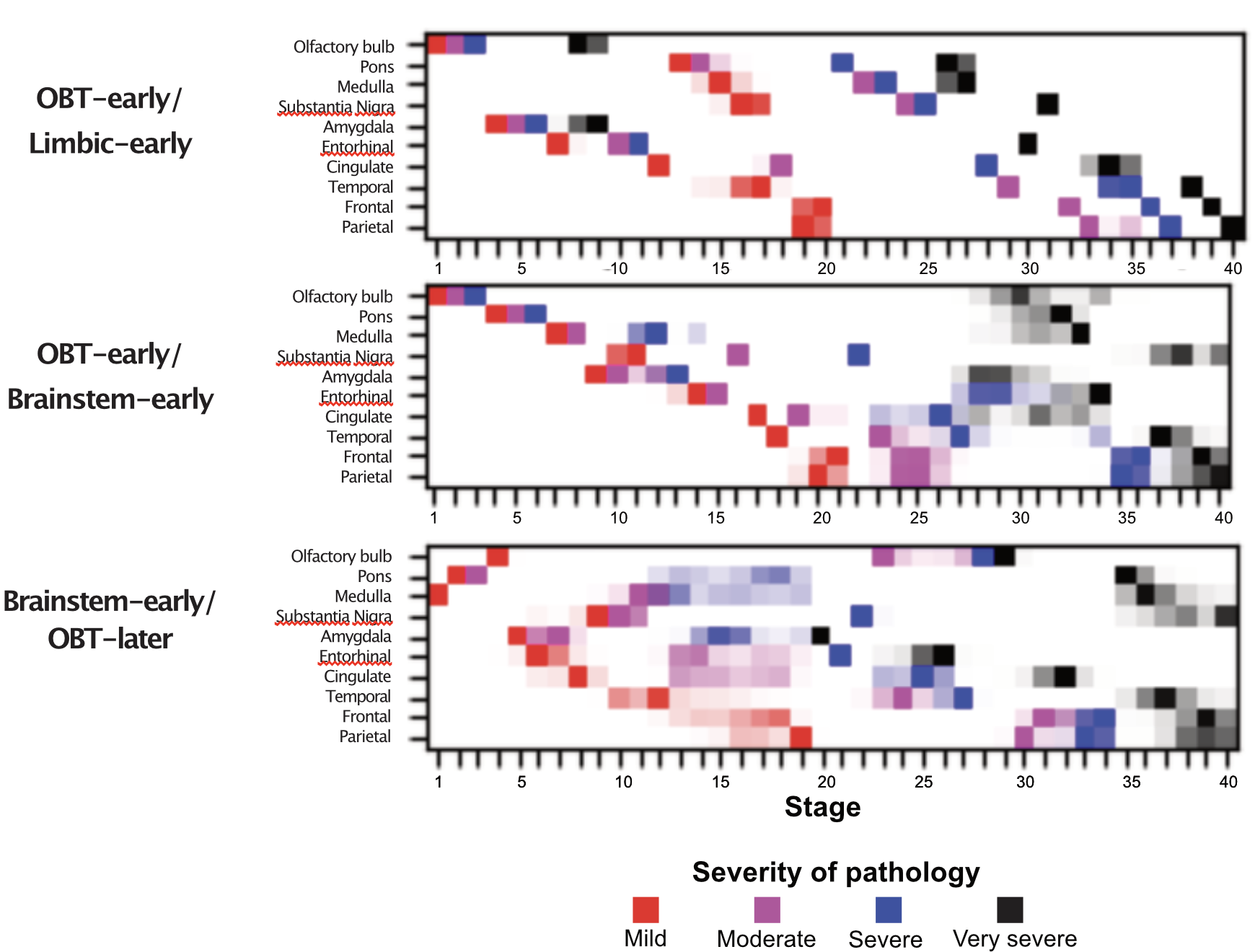

SuStaIn-inferred disease progression patterns of regional Lewy-type α-synuclein pathology in a subset of the BBDP cohort with complete *post-mortem* α-synuclein data (i.e., assessed in all 10 sampled regions). SuStaIn identified 3 subtypes of which the positional variance diagrams are shown, with each box representing the certainty that a brain region has reached a certain level of pathology (red = mild, purple = moderate, blue = severe, black = very severe) at a given SuStaIn or disease progression stage. Darker colors represent more confidence. Trajectories similar to the 3 subtypes in the main sample were identified.

OBT = olfactory bulb and tract; SuStaIn = Subtype and Stage Inference

**Figure S3 Output of the SuStaIn model applied to re-examined OBT data**

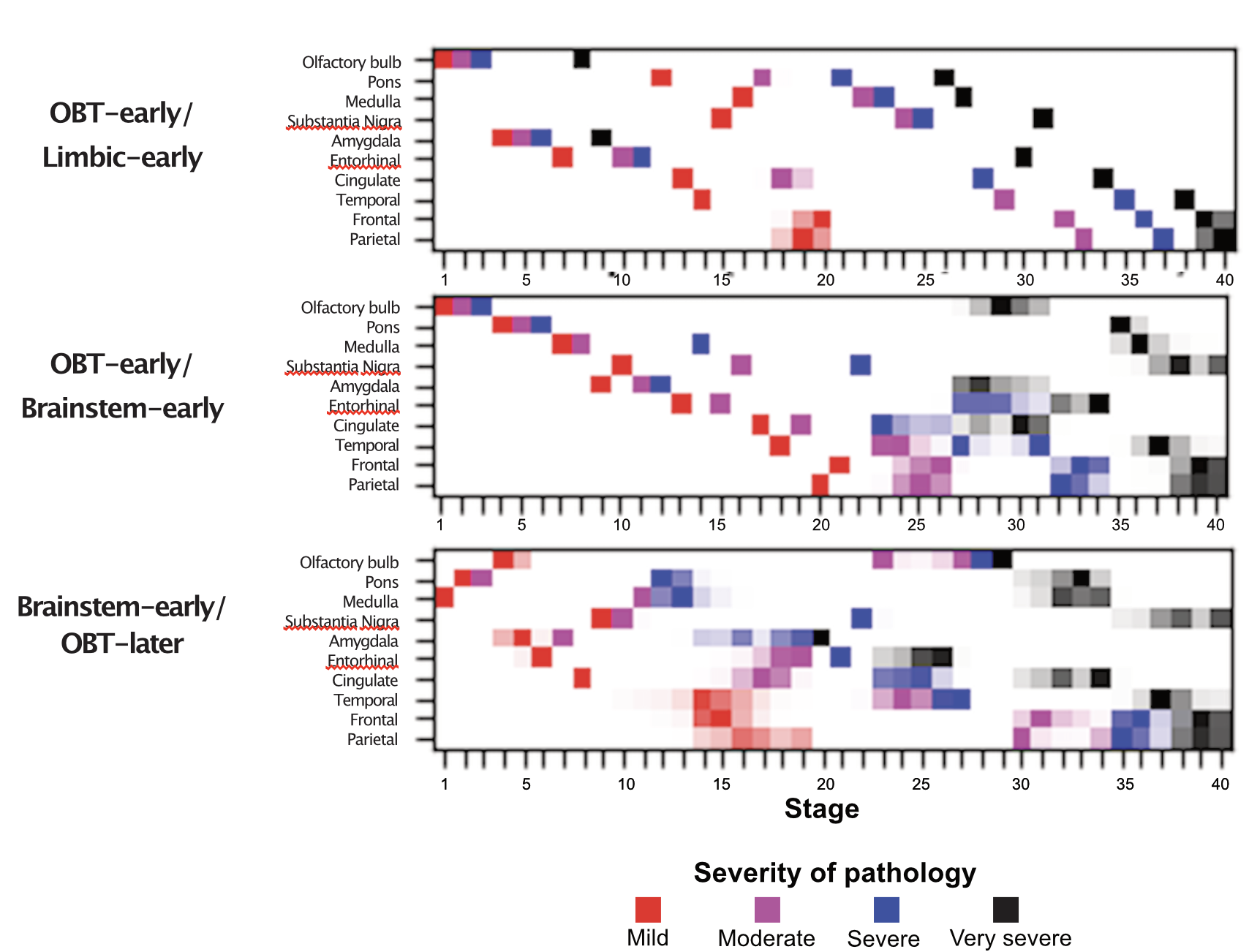

SuStaIn-inferred disease progression patterns of regional Lewy-type α-synuclein pathology in the BBDP cohort with re-examined OBT-stainings in OBT-negative cases of the Brainstem-early/OBT-later subtype. SuStaIn identified 3 subtypes of which the positional variance diagrams are shown, with each box representing the certainty that a brain region has reached a certain level of pathology (red = mild, purple = moderate, blue = severe, black = very severe) at a given SuStaIn or disease progression stage. Darker colors represent more confidence. Trajectories similar to the 3 subtypes in the main dataset were identified.

OBT = olfactory bulb and tract; SuStaIn = Subtype and Stage Inference

**Figure S4. Subtype assignment probability across assigned stage**

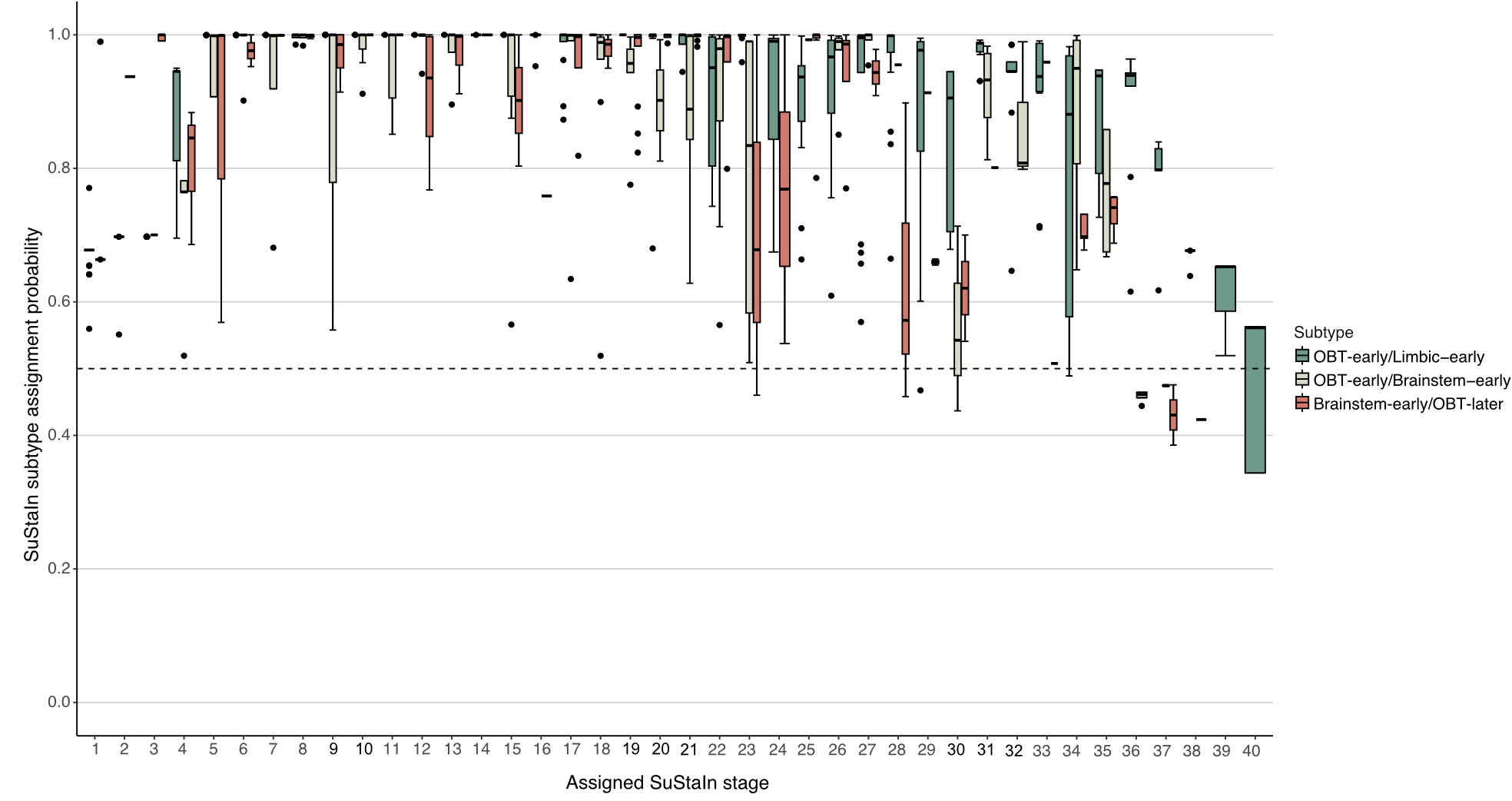

Boxplots show the probability of subtype assignment across assigned stages for the SuStaIn modelling cohort with stage>0. The dashed line represents the cut-off of 50% for high probability. Subtype probability decreases with more advanced stages, reflecting similarity between subtypes.

OBT = olfactory bulb and tract; SuStaIn = Subtype and Stage Inference

**Figure S5. Distribution of stage assignment**

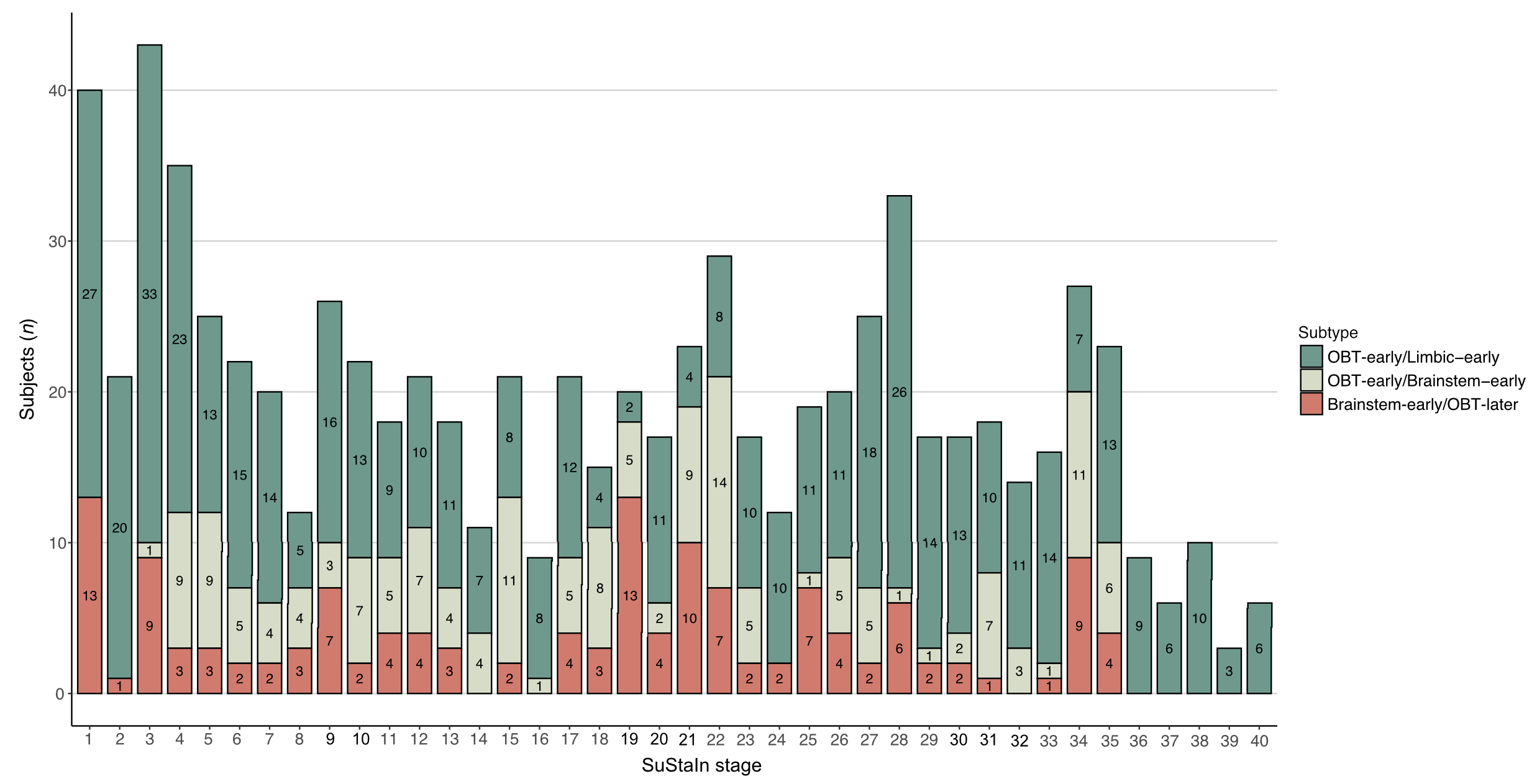

Bar chart shows the number of cases (stage>0, subtype probability>50%) assigned to each SuStaIn stage.

OBT = olfactory bulb and tract; SuStaIn = Subtype and Stage Inference

**Figure S6. Number of subjects in stages of the Unified Staging Scheme for Lewy Body disorders according to SuStaIn stage**

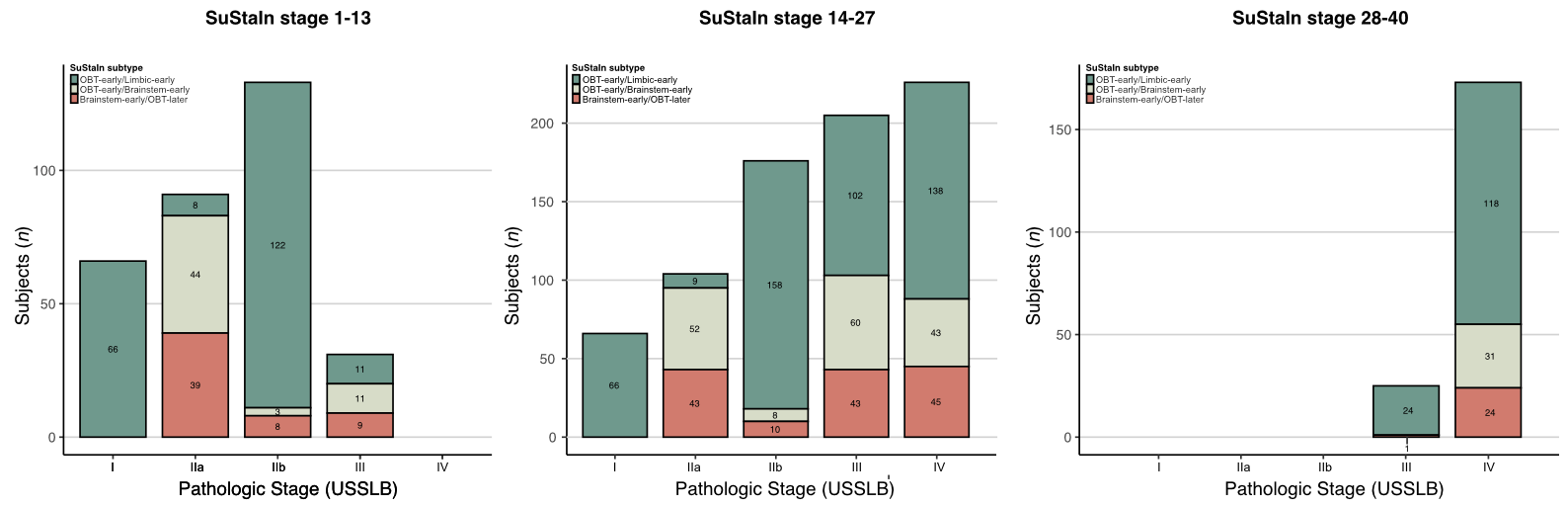

Staging of subjects (*n*=777) according to the Unified Staging Scheme for Lewy Body Disorders (USSLB), colored by SuStaIn subtype. Separate figures are shown for early (1-13), middle (14-27), and late (28-40) SuStaIn stage.

OBT = olfactory bulb and tract; SuStaIn = Subtype and Stage Inference

**Figure S7. Sliding window analysis of LB subtype comparisons**

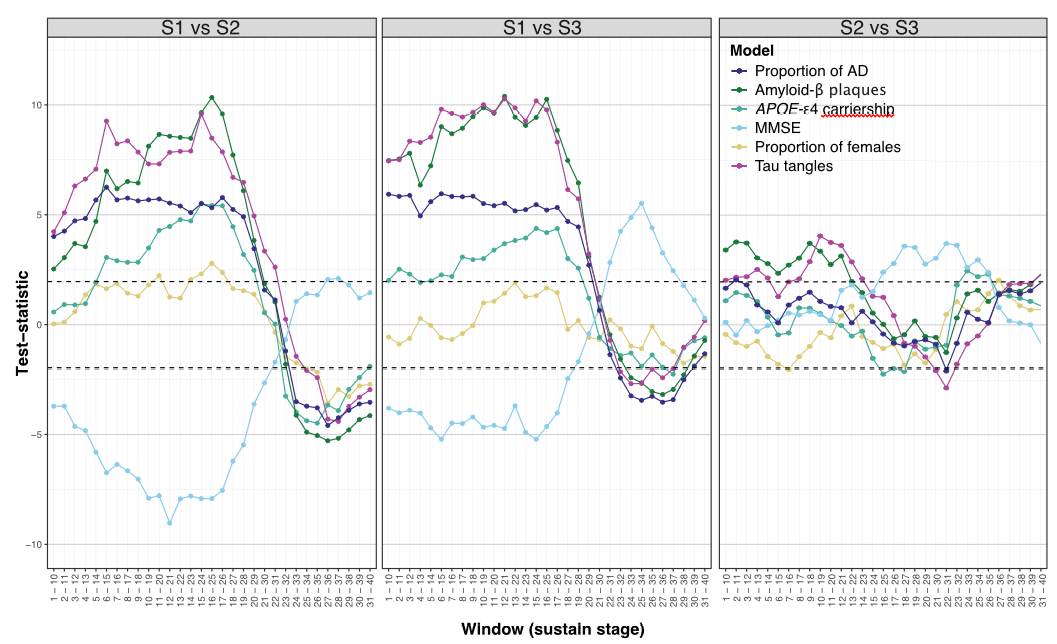

Sliding window analysis (window width = 10, slide = 1) of subtype comparisons on the proportion of subjects with clinicopathological Alzheimer’s disease, plaque burden, *APOE*-ε4 carriership, MMSE scores, the proportion of female subjects, and neurofibrillary burden. Each datapoint represents the test-statistic (*t-*value in case of continuous outcome variable and *z*-statistic in case of categorical outcome variable) of regression models adjusted for age, sex, and SuStaIn stage investigating subtype differences in a specific SuStaIn stage interval.

AD = Alzheimer’s disease; MMSE = Mini Mental State Examination; S1 = OBT-early/Limbic-early; S2 = OBT-early/Brainstem-early; S3 = Brainstem-early/OBT-later.

**Figure S8. Total Lewy body pathology across subtypes with a SuStaIn stage<10**

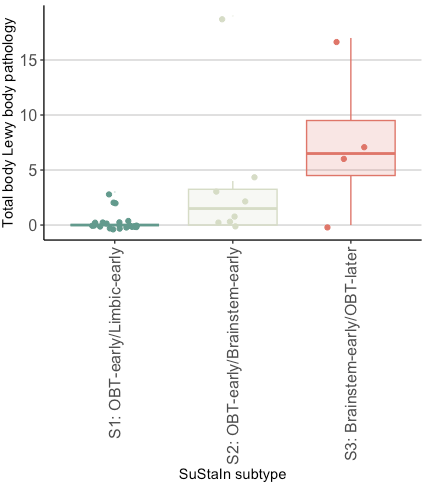

Total pathology scores were computed as the sum of the α-synuclein density scores (0-4) assessed in the cervical, thoracic, lumbar, and sacral spinal cord gray matter, vagus nerve, submandibular gland, and esophagus, with higher scores indicating more severe pathology.

OBT = olfactory bulb and tract
